# Mitohormetic effects of rotenone drastically depend on age

**DOI:** 10.1101/528547

**Authors:** Mario Baumgart, Martino Ugolini, Marco Groth, Matthias Platzer, Alessandro Cellerino

**Author notes:** CORRESPONDENCE TO, Alessandro Cellerino,050 509317, Palazzo della Carovana, terzo piano, stanza 71.

## Abstract

Hormesis refers to a biphasic intensity-dependent response to stressors where low-intensity or short-duration exposure to potentially noxious stressors induces long-lasting adaptations that have positive physiological effects [1]. It was proposed that life-extending interventions, such as calorie restriction, retard aging via their hormetic action [2]. More recently, it was shown that a transient burst of reactive oxygen species is required to induce the effects of calorie restriction [3], physical exercise [4], inhibition of the insulin/IGF-I pathway [5] and metformin [6]. We have previously shown that partial inhibition of complex I of the respiratory chain by a low dose of the poison rotenone (ROT) administered in middle age increases the lifespan and reverts the transcriptomic signature of aging in a vertebrate species [7], an example of a hormetic effect. Here, we asked whether ROT treatment started at young age induces larger effects on life-span and transcriptome.

As an experimental model, we used the short-lived fish *Nothobranchius furzeri* that has become a model of choice to test effects of life-long intervention in vertebrates [8]–[11]. The strain (MZM-0410) used in the present study reaches sexual maturity at about 4 weeks post hatch (wph), has a median lifespan of around 30 wph [7] and expresses conserved transcriptomic signatures associated to vertebrate aging [12]. Fish were treated with ROT at a concentration of 0.1% LC50 (15-20pM) starting at age 5 wph. This concentration of ROT was shown to induce life extension when treatment started at 23 wph [7]. This experiment ran in parallel with the experiment described by Baumgart et al. 2016 [7] in order to exclude differences in the experimental conditions.

Treatment with ROT starting in young age (5 wph) resulted in life-shortening (p=0.0467, Log-rank test, Fig 1). In order to obtain a global evaluation of the effects of ROT treatment, we performed RNA-seq analysis on brain, liver and skin after 4 weeks of treatment (age of analysis 9 wph). We then compared the effects of ROT with the effects of aging on global gene expression visualized as 2D scatterplots of log2(fold changes) in the two conditions. A clear positive correlation was observed in all three tissues, indicating that ROT treatment at young ages shifts the transcriptome towards a pattern that is more typical of older ages. We then compared the effects of ROT at the two different ages and observed a strong negative correlation in all three tissues, indicating that the differentially expressed genes are more likely to be regulated in opposite directions by ROT in young and old animals.

**Figure 1.**
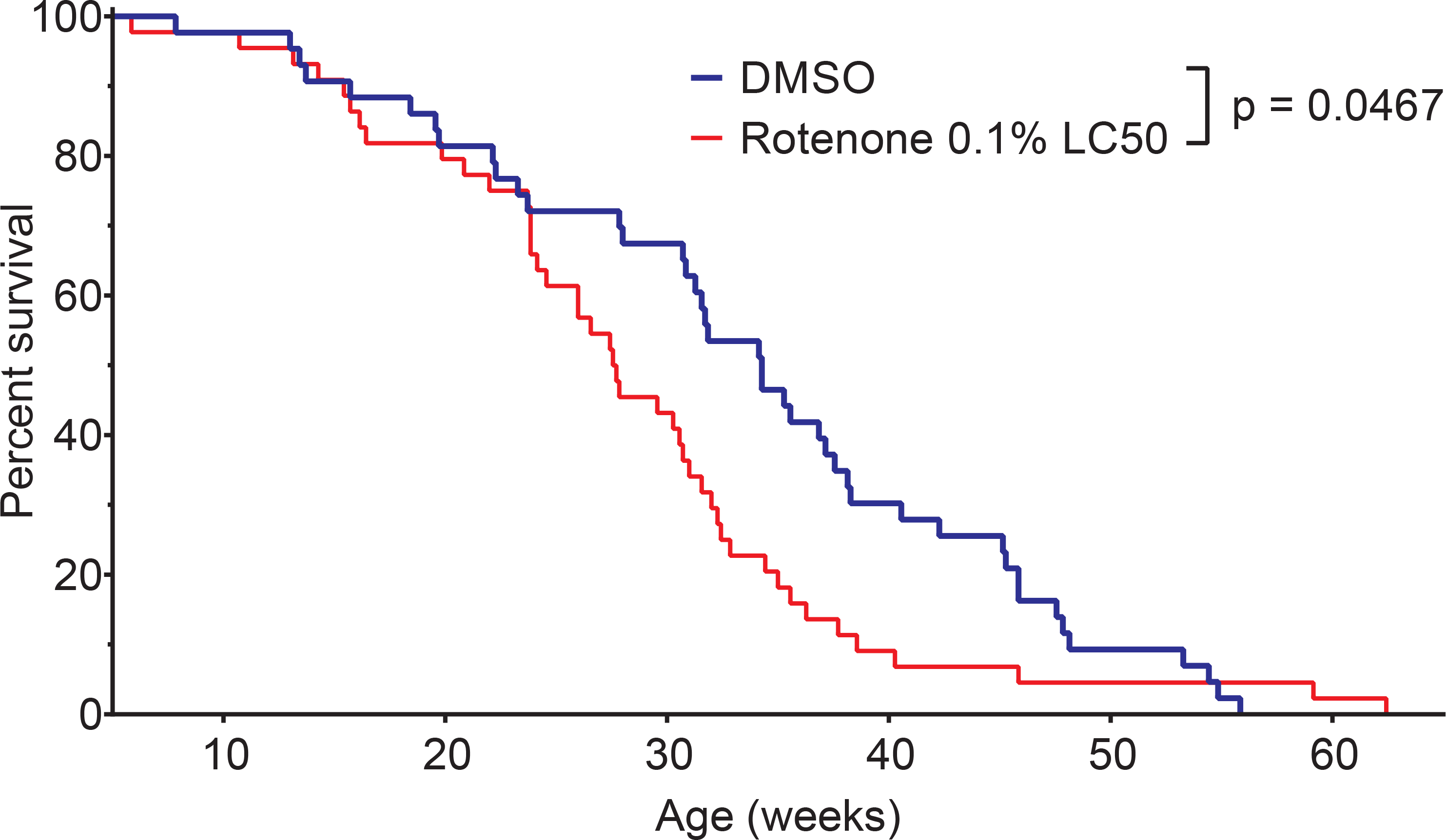
Effects of rotenone on lifespan. Survival of rotenone- and vehicle-treated *N. furzeri*; Rotenone 0.1% LC50 (n =44, median survival = 27,6 weeks, red line) and DMSO (n =43, median survival = 34,3 weeks, blue line); p values, Log-rank test.

We then performed a meta-analysis to identify GO categories that are enriched in genes regulated in opposite directions by ROT depending on age. Genes up-regulated in young and down-regulated in old fish were enriched in several different categories (Fig 2G), genes up-regulated in old and down-regulated in young fish provided a more structured picture and were enriched for terms related to biosynthesis and trafficking of RNA, proteins and ribonucleoprotein complexes as well as stress responses such as ER stress and p53 pathway (Fig 2H).

**Figure 2.**
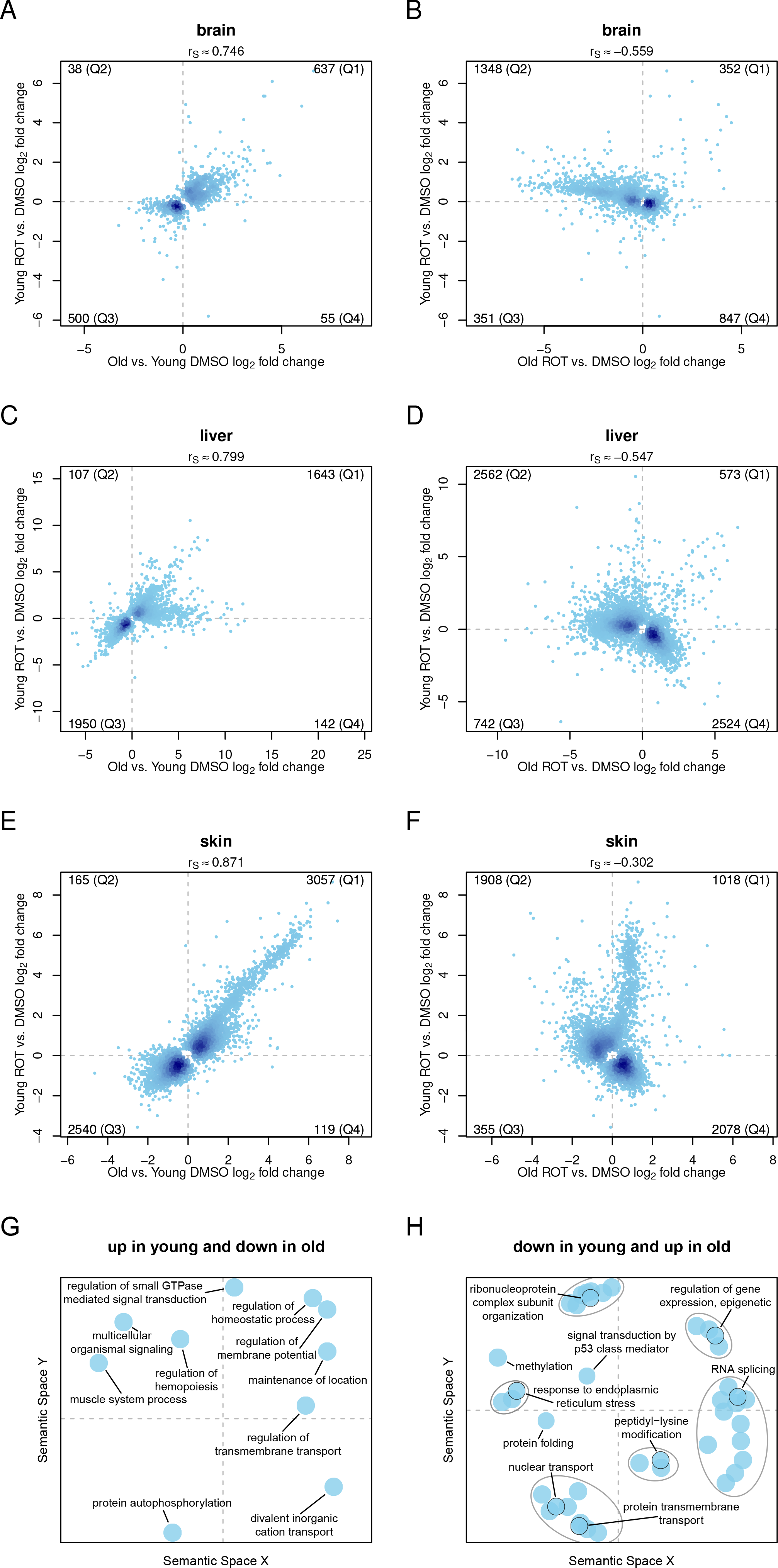
Effects of rotenone (ROT) on global gene expression visualized as scatterplots. Each point corresponds to one gene and darker colours correspond to overlap of points. The number of genes plotted in each graph correspond to the union of the genes differentially expressed in either condition. The numbers in the four quadrants indicate the number of genes contained in the quadrant. (A-F) Effects of 4 weeks treatment of ROT on brain (A,B), liver (C,D) and skin (E,F). In (A),(C) and (E) the effects of ROT treatment in young are compared with those of aging. The X-axis corresponds to the effects of age quantified as log2 of the ratio in the expression levels [old(27 weeks) controls]/[young(9 weeks) controls] and the Y-axis corresponds to the effects of ROT in young animals quantified as log2 of the ratio in the expression levels treated/controls. Number of genes whose regulation is plotted in (A),(C) and (E) is 1230, 3842 and 5881, respectively. In (B),(D),(F) the effects of ROT at two different ages is compared. The X-axis corresponds to the effects of ROT in old quantified as log2 of the ratio in the expression levels [old(27 weeks) ROT]/[old(27 weeks) controls] and the X-axis corresponds to the effects of ROT in young quantified as log2 of the ratio in the expression levels [young (9 weeks) ROT]/[young (9 weeks) controls]. Number of genes whose regulation is plotted in (B),(D) and (F) is 2898, 6401 and 5359, respectively. (G,H) GO overrepresentation analysis for genes that are (G) up-regulated in old and down-regulated in young or (H) up-regulated in young and down-regulated in old. Graphs were generated by REVIGO and represent multidimensional scaling using a measure of semantic dissimilarity. Each dot represents a single GO category that is significantly overrepresented (FDR < 0.05). The distances between dots are inversely related to the similarity of the different categories. Ellipses in (H) represent clusters of highly similar GO categories.

In summary, our work shows that a mild increase in mitochondrial ROS production has radical different outcomes in the context of an old and a young organism. A similar age-dependent antagonistic action was previously reported for what concerns the effects of mitochondrial catalase overexpression on heart protein turnover [13] and appears as a “reverse” antagonistic pleiotropism [14].

## MATERIAL AND METHODS

Materials and Methods are available in the Supplements.

## DATA AVILABILITY

For this work a subset of the dataset GSE66712 (already used in [18]) and the three datasets GSE103804, GSE103809 and GSE103819, have been analysed. Please refer to supplementary table 1 for further details.

## AUTHOR CONTRIBUTION

MB, MP and AC conceived and designed the study; MB performed the *N. furzeri* experiments; MG performed the RNA-seq; MU performed data analysis; AC wrote the manuscript with contributions from all other authors.

## ACKNOLEDGEMENTS

This work was partially supported by the German Ministry for Education and Research (JenAge; BMBF support code 0315581A) and by intramural grant of Scuola Normale Superiore.

## SUPPLEMENTARY MATERIALS

### Materials and Methods

**Table S1.** Overview of GEO data sets and files

| 9 weeks, DMSO |  |  |  |  |  |
| --- | --- | --- | --- | --- | --- |
| Brain (GSE66712) |  | Liver (GSE66712) |  | Skin (GSE66712) |  |
| GSM1630266 | MC1_1 | GSM1630267 | MC1_3 | GSM1630268 | MC1_5 |
| GSM1630269 | MC2_1 | GSM1630270 | MC2_3 | GSM1630271 | MC2_5 |
| GSM1630272 | MC3_1 | GSM1630273 | MC3_3 | GSM1630274 | MC3_5 |
| GSM1630275 | MC4_1 | GSM1630276 | MC4_3 | GSM1630277 | MC4_5 |
| GSM1630278 | MC5_1 | GSM1630279 | MC5_3 | GSM1630280 | MC5_5 |
| 27 weeks, DMSO |  |  |  |  |  |
| Brain (GSE66712) |  | Liver (GSE66712) |  | Skin (GSE66712) |  |
| GSM1630284 | MC6_1 | GSM1630285 | MC6_3 | GSM1630286 | MC6_5 |
| GSM1630287 | MC7_1 | GSM1630288 | MC7_3 | GSM1630289 | MC7_5 |
| GSM1630290 | MC8_1 | GSM1630291 | MC8_3 | GSM1630292 | MC8_5 |
| GSM1630293 | MC9_1 | GSM1630294 | MC9_3 | GSM1630295 | MC9_5 |
| GSM1630281 | MC10_1 | GSM1630282 | MC10_3 | GSM1630283 | MC10_5 |
| 9 weeks, Rotenone 0.1% LC50 |  |  |  |  |  |
| Brain (GSE103804) |  | Liver (GSE103809) |  | Skin (GSE103819) |  |
| GSM3403526 | MR2_1 | GSM3334273 | MR2_3 | GSM3334278 | MR2_5 |
| GSM3403527 | MR3_1 | GSM3334274 | MR3_3 | GSM3334279 | MR3_5 |
| GSM3403528 | MR6_1 | GSM3334275 | MR6_3 | GSM3334280 | MR6_5 |
| GSM3403529 | MR7_1 | GSM3334276 | MR7_3 | GSM3334281 | MR7_5 |
| GSM3403525 | MR10_1 | GSM3334272 | MR10_3 | GSM3334277 | MR10_5 |
| 27 weeks, Rotenone 0.1% LC50 |  |  |  |  |  |
| Brain (GSE66712) |  | Liver (GSE66712) |  | Skin (GSE66712) |  |
| GSM1630296 | MR26_1 | GSM1630297 | MR26_3 | GSM1630298 | MR26_5 |
| GSM1630299 | MR27_1 | GSM1630300 | MR27_3 | GSM1630301 | MR27_5 |
| GSM1630302 | MR28_1 | GSM1630303 | MR28_3 | GSM1630304 | MR28_5 |
| GSM1630305 | MR29_1 | GSM1630306 | MR29_3 | GSM1630307 | MR29_5 |
| GSM1630308 | MR30_1 | GSM1630309 | MR30_3 | GSM1630310 | MR30_5 |

## MATERIALS AND METHODS

### Fish maintenance

In this paper, the *N. furzeri* strain MZM-0410 [1] was studied. Animal maintenance was performed as described [1], [2]. The same setup was used for all experiments and animals were raised on a rolling base. To avoid effects of circadian rhythms and feeding, animals were always sacrificed at 10 a.m. in fasted state. For tissue preparation, fish were euthanized with MS-222 and cooled on crushed ice. The protocols of animal maintenance and experimentation were approved by the local authority in the State of Thuringia (Veterinaer- und Lebensmittelueberwachungsamt, Rotenone treatment: reference number 22-2684-04-03-003/13).

### Rotenone treatment

Rotenone is a natural toxin produced by several tropical plants. It is a highly specific metabolic poison that affects cellular aerobic respiration, blocking mitochondrial electron transport by inhibiting NADH ubiquinone [3]. Rotenone is highly toxic to fish, with median lethal concentration in water (LC50) commonly between 5 and 250 nM. Individual species sensitivities vary widely, with salmonids most sensitive and goldfish, carp and bullhead catfish least so [4]. Reasons for such variation in sensitivity may reside in differences in the levels of liver enzymes responsible for the chemical breakdown or detoxification of Rotenone [4].

For the determination of Rotenone lethality in *N. furzeri*, Rotenone was dissolved in DMSO at a working concentration of 1 mM. In groups of 10 fish (tank size of 5 L and standard water conditions) we tested lethality within 96 hours as described in Baumgart et al., 2016. LC50 96h for *N. furzeri* was determined to be 20 nM.

*N. furzeri* were housed in groups of 15 animals in 40 L tanks with standard water conditions. Rotenone dissolved in DMSO or equal amounts of pure DMSO was added to final concentrations of 20 pM. Treatment started when the fish reached the age of 5 weeks (young group) or 23 weeks (old group). Water quality was maintained by 30% water change twice per week and fresh Rotenone and DMSO was added to the tanks. After 4 weeks, 5 animals of each group were sacrificed and RNA was extracted from brain, liver and skin tissues. The remaining animals were treated until end of life and each death was recorded to plot survival curves. GraphPad Prism 7.01 was used to plot survival curves and to calculate p-value (Log-rank test).

### RNA extraction

Tissue samples were transferred into 2 ml tubes with 1 ml cooled QIAzol (Qiagen, Hilden, Germany) and one 5 mm stainless steel bead (Qiagen) was added. Homogenization was performed using a TissueLyzer II (Qiagen) at 30 Hz for 3x 1 min. After incubation for 5 min at room temperature 200 μl chloroform was added. The tube was shaken for 15 s and incubated for 3 min at room temperature. Phase separation was achieved by centrifugation at 12,000x g for 20 min at 4°C. The aqueous phase was transferred into a fresh cup and 10 μg of Glycogen (Invitrogen, Darmstadt, Germany), 0.16x volume NaAc (2 M; pH 4.2) and 1.1x volume isopropanol were added, mixed and incubated for 10 min at room temperature. The RNA was precipitated by a centrifugation step with 12,000x g at 4°C for 20 min. The supernatant was removed and the pellet was washed with 80% Ethanol twice and air dried for 10 min. The RNA was resuspended in 20 μl nuclease-free water by pipetting up and down, followed by incubation at 65°C for 5 min. The RNA was quantified with a NanoDrop 1000 (PeqLab, Erlangen, Germany) and stored at - 80°C until use.

### RNA-sequencing

Library preparation and sequencing was done using Illumina’s NGS platform [5]. In detail, total RNA was quality checked using an Agilent Bioanalyzer 2100 and Agilent RNA 6000 Nano Kit (Agilent Technologies). RNA integrity numbers range from 7.7 to 10 with an average of 9. One μg of total RNA was used for library preparation using Illumina’s TruSeq RNA sample prep kit v2 following the manufacturer’s instruction. A step of purification of the polyA+ fraction was included and this prevented the analysis of ribosomal RNAs. Libraries were quality checked and quantified using Agilent Bioanalyzer 2100 and Agilent DNA 7500 kit. Libraries were sequenced using an Illumina HiSeq2500 in high-output, 50bp single-read mode in pools of four or five per lane. Sequencing chemistry v3 was used.

Read data were extracted in FastQ format using the Illumina supported tool bcl2fastq v1.8.4. Sequencing resulted in around 50mio or 39mio reads per sample when pooling four or five samples per lane, respectively.

### RNA-seq data analysis

Raw sequencing data were received in FASTQ format. Read mapping was performed using Tophat 2.0.6 [6] and a pre-release version of the N. furzeri genome assembly PRJEB5837 which corresponds to the scaffolds with the accessions LN600428-LN606439. The resulting SAM alignment files were processed using featureCounts v1.4.3-p1 [7] and the respective GTF gene annotation file.

### Identification of DEGs

Please note that all analyses were performed using R (version 3.5.1) [8] and R-Studio (version 1.1.456) [9]. Starting from the raw counts the log_2_FCs for all genes were estimated – for each tissue separately – in the three conditions “old vs. young DMSO treated animals”, “Rotenone vs. DMSO treated young animals” and “Rotenone vs. DMSO treated old animals” using DESeq2 (version 1.20.0) [10].

The effect of two different conditions on the gene expression profiles were visualized by plotting on the Cartesian plane the log_2_FC in the two conditions. Please note that only the genes that are differentially expressed (adjusted p-value<0.05) in at least one of the two conditions were shown. For each plot the Spearman correlation coefficient r_S_ [11] was estimated (p-value<2.2e-16) and its value – rounded to 3 decimal places – shown in the plots. The number of genes present in each quadrant is also shown in the corresponding corners.

### Gene Enrichment Analysis

For each quadrant of the plots described in the previous section an over representation analysis (ORA) was performed with the R package of WebGestalt (version 0.1.1) [12] using the human orthologs of the genes contained in them. The categories of the GO database [13], [14] that contained at least 5 genes were considered, and the p-values were corrected for multiple comparisons through the FDR method described in [15].

The categories that are enriched in a quadrant, independently from the considered tissue, were identified through meta-analysis using Fisher’s method [16], and the obtained p-values were corrected for multiple comparisons through the FDR method described in [15]. Finally, the categories with adjusted p-value<0.05 were visualized using REVIGO [17], a method that removes redundant categories and visualizes the remaining representative ones based on their sematic similarity.

## REFERENCES

[1] E. J. Calabrese and L. A. Baldwin, “Defining hormesis,” Hum. Exp. Toxicol., vol. 21, no. 2, pp. 91–97, Feb. 2002.

[2] E. J. Masoro, “Hormesis and the antiaging action of dietary restriction.,” Exp. Gerontol., vol. 33, no. 1–2, pp. 61–6.

[3] T. J. Schulz, K. Zarse, A. Voigt, N. Urban, M. Birringer, and M. Ristow, “Glucose Restriction Extends Caenorhabditis elegans Life Span by Inducing Mitochondrial Respiration and Increasing Oxidative Stress,” Cell Metab., vol. 6, no. 4, pp. 280–293, Oct. 2007.

[4] M. Ristow et al., “Antioxidants prevent health-promoting effects of physical exercise in humans,” Proc. Natl. Acad. Sci., vol. 106, no. 21, pp. 8665–8670, May. 2009.

[5] K. Zarse et al., “Impaired insulin/IGF1 signaling extends life span by promoting mitochondrial L-proline catabolism to induce a transient ROS signal.,” Cell Metab., vol. 15, no. 4, pp. 451–65, Apr. 2012.

[6] W. De Haes et al., “Metformin promotes lifespan through mitohormesis via the peroxiredoxin PRDX-2.,” Proc. Natl. Acad. Sci. U. S. A., vol. 111, no. 24, pp. E2501–9, Jun. 2014.

[7] M. Baumgart et al., “Longitudinal RNA-Seq Analysis of Vertebrate Aging Identifies Mitochondrial Complex I as a Small-Molecule-Sensitive Modifier of Lifespan.,” Cell Syst., vol. 2, no. 2, pp. 122–32, Feb. 2016.

[8] A. Cellerino, D. R. Valenzano, and M. Reichard, “From the bush to the bench: the annual Nothobranchius fishes as a new model system in biology.,” Biol. Rev. Camb. Philos. Soc., vol. 91, no. 2, pp. 511–33, May. 2016.

[9] Y. Kim, H. G. Nam, and D. R. Valenzano, “The short-lived African turquoise killifish: an emerging experimental model for ageing.,” Dis. Model. Mech., vol. 9, no. 2, pp. 115–29, Feb. 2016.

[10] M. Platzer and C. Englert, “Nothobranchius furzeri : A Model for Aging Research and More,” Trends Genet., vol. 32, no. 9, pp. 543–552, Sep. 2016.

[11] C.-K. Hu and A. Brunet, “The African turquoise killifish: A research organism to study vertebrate aging and diapause.,” Aging Cell, vol. 17, no. 3, p. e12757, Jun. 2018.

[12] P. Aramillo Irizar et al., “Transcriptomic alterations during ageing reflect the shift from cancer to degenerative diseases in the elderly.,” Nat. Commun., vol. 9, no. 1, p. 327, Dec. 2018.

[13] N. Basisty et al., “Mitochondrial-targeted catalase is good for the old mouse proteome, but not for the young: ‘reverse’ antagonistic pleiotropy?,” Aging Cell, vol. 15, no. 4, pp. 634–645, Aug. 2016.

[14] G. C. Williams, “Pleiotropy, natural selection and the evolution of senescence,” Evolution (N. Y)., vol. 11, no. 4, pp. 398–411, 1957.

## REFERENCES

[1] E. Terzibasi et al., “Large differences in aging phenotype between strains of the short-lived annual fish Nothobranchius furzeri.,” PLoS One, vol. 3, no. 12, p. e3866, Dec. 2008.

[2] E. Terzibasi, C. Lefrançois, P. Domenici, N. Hartmann, M. Graf, and A. Cellerino, “Effects of dietary restriction on mortality and age-related phenotypes in the short-lived fish Nothobranchius furzeri.,” Aging Cell, vol. 8, no. 2, pp. 88–99, Apr. 2009.

[3] T. P. Singer and R. R. Ramsay, “The reaction sites of rotenone and ubiquinone with mitochondrial NADH dehydrogenase.,” Biochim. Biophys. Acta, vol. 1187, no. 2, pp. 198–202, Aug. 1994.

[4] N. Ling, “Rotenone - a review of its toxicity and use for fisheries management,” Sci. Conserv., 2003.

[5] D. R. Bentley et al., “Accurate whole human genome sequencing using reversible terminator chemistry,” Nature, vol. 456, no. 7218, pp. 53–59, Nov. 2008.

[6] D. Kim, G. Pertea, C. Trapnell, H. Pimentel, R. Kelley, and S. L. Salzberg, “TopHat2: accurate alignment of transcriptomes in the presence of insertions, deletions and gene fusions.,” Genome Biol., vol. 14, no. 4, p. R36, Apr. 2013.

[7] Y. Liao, G. K. Smyth, and W. Shi, “featureCounts: an efficient general purpose program for assigning sequence reads to genomic features.,” Bioinformatics, vol. 30, no. 7, pp. 923–30, Apr. 2014.

[8] R. D. C. Team and R. R Development Core Team, “R: A Language and Environment for Statistical Computing,” R Found. Stat. Comput., 2016.

[9] RStudio, “RStudio: Integrated development environment for R (Version 0.97.311),” J. Wildl. Manage. 2011.

[10] M. I. Love, W. Huber, and S. Anders, “Moderated estimation of fold change and dispersion for RNA-seq data with DESeq2,” Genome Biol., vol. 15, no. 12, p. 550, Dec. 2014.

[11] C. Spearman, “The Proof and Measurement of Association between Two Things,” Am. J. Psychol., vol. 100, no. 3/4, p. 441, 1987.

[12] J. Wang, D. Duncan, Z. Shi, and B. Zhang, “WEB-based GEne SeT AnaLysis Toolkit (WebGestalt): update 2013,” Nucleic Acids Res., vol. 41, no. W1, pp. W77–W83, Jul. 2013.

[13] The Gene Ontology Consortium, “Expansion of the Gene Ontology knowledgebase and resources,” Nucleic Acids Res., vol. 45, no. D1, pp. D331–D338, Jan. 2017.

[14] M. Ashburner et al., “Gene Ontology: tool for the unification of biology,” Nat. Genet., vol. 25, no. 1, pp. 25–29, May. 2000.

[15] Y. Benjamini and Y. Hochberg, “Controlling the False Discovery Rate: A Practical and Powerful Approach to Multiple Testing,” Journal of the Royal Statistical Society. Series B (Methodological), vol. 57. WileyRoyal Statistical Society, pp. 289–300, 1995.

[16] R. Fisher, Statistical Methods for Research Workers, 1st ed. Edinburgh: Oliver & Boyd, 1925.

[17] F. Supek, M. Bošnjak, N. Škunca, and T. Šmuc, “REVIGO summarizes and visualizes long lists of gene ontology terms.,” PLoS One, vol. 6, no. 7, p. e21800, Jul. 2011.

[18] M. Baumgart et al., “Longitudinal RNA-Seq Analysis of Vertebrate Aging Identifies Mitochondrial Complex I as a Small-Molecule-Sensitive Modifier of Lifespan.,” Cell Syst., vol. 2, no. 2, pp. 122–32, Feb. 2016.

